# Development of African Swine Fever Virus Pseudotype Systems for Ba71v and Georgia 2007 strains

**DOI:** 10.64898/2026.11.12.765316

**Authors:** George L Hooper, Christopher L Netherton, Edward Wright

## Abstract

African swine fever virus (ASFV) is a large, complex DNA virus that causes a severe haemorrhagic disease in domestic swine and Eurasian wild boar. Due to its high pathogenicity, research involving ASFV is restricted to high-containment laboratories, which significantly limits investigations into its biology, host interactions and the development of effective control measures. To overcome these limitations, this study utilised pseudotyped virus (PV) technology to identify ASFV proteins involved in cell entry. Candidate proteins were identified through bioinformatic analysis and a review of the published literature, incorporated into PVs, and subsequently assessed for their ability to facilitate cellular entry. This approach identified a combination of three ASFV proteins that, when combined, resulted in the most efficient cell entry. This work represents the successful generation of the first functional ASF pseudotype system for both Ba71v and Georgia 2007 strains, enabling the safe study of ASFV entry mechanisms under containment level 2.

## Introduction

ASFV causes a severe haemorrhagic disease in domestic swine and Eurasian wild boar, with virulent strains exhibiting exceptionally high case fatality rates of up to 100% in naïve populations (Blome *et al*., 2020). The current panzootic began with the emergence of the disease in Georgia in 2007, since then ASFV has spread rapidly through the Caucasus region into the Russian Federation, Eastern Europe and most recently into numerous Asian countries. This includes the spread of the virus, in 2018, to the world’s largest producer and consumer of pork, China (Zhou *et al*., 2018).

The high mortality of acute ASFV infection and the lack of safe and effective vaccines or antivirals has enabled the virus to spread rapidly, causing the deaths of millions of wild and domestic pigs, massive economic losses, crippling the pig industry and threatening biodiversity (FAO-Situation-Report, 2019, Smith *et al*., 2019, Zhou *et al*., 2018, Lu *et al*., 2020).

Due to its high pathogenicity, research involving ASFV is restricted to high-containment (SAPO4 in the United Kingdom) laboratories, which significantly limits investigations into its biology, host interactions and the development of effective control measures. To overcome the barriers of high-containment, tools such as pseudotyped viruses (PV) can be implemented. PV are replication defective viral particles that comprise of the structural core and non-structural proteins of one virus, such as the commonly used, vesicular stomatitis virus (VSV) or human immunodeficiency virus (HIV), which have had key proteins involved in viral replication deleted. Expressing the envelope protein(s) of the of virus of interest in the same cells as the core proteins leads to those envelope protein(s) being incorporated into the PV (Naldini *et al*., 1996, Zufferey *et al*., 1997, Dull *et al*., 1998, Burns *et al*., 1993). Thus, a PV comprises the core of one virus and the viral envelope protein(s) (VEP) of another, and enables the role of proteins in early events in the virus’ life cycle to be tested without requiring infectious virus, and in the case of viruses such as ASFV without high containment laboratories.

The ASFV’s genome ranges from 170-194 kb, containing 151-200 open reading frames (Cackett *et al*., 2022), with many of the proteins uncharacterised or poorly characterised. The ASFV genome resides within a nucleoid, that is encased within a core shell, referred to as the inner capsid. The core is then surrounded by an inner envelope, derived from the host cell’s endoplasmic reticulum, obtained during virus morphogenesis in perinuclear viral factories (Andres *et al*., 1998, Rouiller *et al*., 1998, Salas and Andrés, 2013). The inner envelope is encased by another capsid, formed from the major capsid protein (MCP) p72, encoded by the gene *B646L*, and four minor capsid proteins: pM1249L, p17, p49 and pH240R, and is referred to as the outer capsid (Alejo *et al*., 2018, Andrés *et al*., 2020, Wang *et al*., 2019). A pleomorphic outer envelope, obtained from the plasma membrane during the budding process, surrounds the virion, which does not conform to the icosahedral morphology of the capsids and inner envelope, and is not required for infectivity (Alejo *et al*., 2018, Andrés *et al*., 2020, Wang *et al*., 2019, Andres *et al*., 2001).

To-date only one protein, pEP402R (also known as CD2v), a homologue of the human T-lymphocyte surface antigen CD2, has been definitively localised to the outer envelope (Alejo *et al*., 2018). Other potential membrane proteins that have been implicated in entry, which are discussed in detail in (Hooper *et al*., 2024), are p30 (*CP204L*), p54 (*E183L*), pE199L, pE248R, and p12 (*O61R*), which are reported to be located on the internal membrane. Some of these proteins have been shown to interact with host factors, such as CD163, Siglec1, CD1d, and NPC1.

With no known VEP and a large number of poorly characterised proteins generating a ASFV PV is challenging. Here we use a combination of literature sources, bioinformatics and experiments with PV displaying different ASFV proteins to identify key proteins involved in entry and generate an ASFV pseudotype virus system to enable research in low containment facilities.

## Results

### Candidate protein selection

As the protein(s) mediating ASFV cell entry remain unknown, candidate proteins were chosen based on several criteria. First, evidence in the literature to support an involvement in cell entry. This led to the selection of 6 proteins: *CP204L* (p30), *E183L* (p54), *E199l* (pE199L), *E248R* (pE248R), *EP402R* (pEP402R) and *O61R* (p12). Second, whether the protein was predicted to contain a transmembrane domain or a signal peptide. Capsid proteins were excluded as structural proteins are unable to be incorporated into enveloped PV. Further to this, only proteins that had been definitively identified in highly purified extracellular ASFV particles by mass spectrometry were selected (Alejo *et al*., 2018). These criteria led to the selection of a further 13 candidate proteins: pB117L, pB169L, pC257L, pCP123L, pD117L, pEP84R, pEP152R, pEP153R, pF49L, pH108R, pI177L, pKP177R. Although it did not meet all the criteria, X69R was included as a candidate protein because it is an uncharacterised transmembrane protein that is not in the MGF. This gave a total of 19 proteins to take forward for further analysis.

### 3D structural predictions of ASFV candidate VEP

3D structural models were generated for the selected Georgia and Ba71v proteins using Alphafold V3 (Jumper *et al*., 2021, Varadi *et al*., 2024). At the time this work was initiated, only AF2 models for proteins from the Georgia strain of ASFV had been published. Therefore, AF2 models for the 19 candidate proteins were generated for the Ba71V strain. Following the release of AF3, structural models for all 19 candidate proteins from both the Georgia and Ba71V strains were subsequently generated, although there was no major differences in the predicted structures (data not shown). More recently, the Viro3D (Litvin *et al*., 2025) resource was launched, providing predicted structural proteomes for multiple ASFV strains.

### Homology search using 3D structural models

To elucidate the functions of candidate proteins we also searched for structural homologues, first, the model with the highest confidence score for each protein was uploaded to the Foldseek server (van Kempen *et al*., 2024). Using the e-value to filter hits, a cut off of 1 × 10^-3^ was applied to infer there was significant structural homology. Next models of both the Ba71v and Georgia strains were run through Foldseek, however, there was no significant difference in the results. Four of the 19 candidate proteins showed significant homology to other proteins namely, pEP402R, pE248R, pKP177L and pEP153R.

#### *EP402R* (pEP402R)

The lowest e-value hit for pEP402R from the PDB100 was 9.82 × 10^-13^, which showed significant homology with the crystal structure of the extracellular domain of the human cell adhesion molecule CD2 (Bodian *et al*., 1994). Both models produced TM-scores of 0.49 and RMSD values of 6.66 and 6.32 for the Ba71v and Georgia models respectively. This was not surprising given EP*402R* has already been identified as a CD2 homologue. However, the top Foldseek hit from the PDB100 with a e-value of 5.64 × 10^-12^ was with the crystal structure of the CD2 binding domain of CD58 (Ikemizu *et al*., 1999), otherwise known as lymphocyte function-associated antigen 3 (LFA-3), the binding partner of CD2. This hit produced TM-scores of 0.58 and 0.56 and RMSD values of 5 and 4.94 for Ba71v and Georgia models respectively.

#### E*248R* (pE248R)

The only significant hit from the foldseek search of pE248R was from the Georgia strain model, with a e-value of 7.28 × 10^-3^. The hit was to an Alphafold model of apolipoprotein Eb (AF-O42364-F1-v4) from the Alphafold protein structure database, a protein involved in lipid transport and metabolism, which also acts as a ligand for the LDL receptor family (Mahley and Rall, 2000, Weisgraber, 1994). Whilst there is some structural homology between the models, a TM-score of 0.4 and RMSD of 10.18 also shows there is a great deal of divergence between the models and thus, are unlikely to be homologues.

#### *KP177L* (pKP177L)

The hit with the lowest e-value for the Georgia strain was Low-Density Lipoprotein Receptor-Related Protein 11 (LRP11) (AF-Q86VZ4-F1-v4), a receptor from the LDL family of receptors (Sun *et al*., 2024), with an e-value of 7.47 × 10⁻³. For the Ba71v strain, the top hit was GH10711 (AF-Q9Y110-F1-model_v4), a membrane protein from *Drosophila melanogaster* with an unknown function.

A notable similarity between proteins from both strains was that many of the top results in Foldseek contained either a MANSC domain or the structurally related apple domain. Both LRP11 and GH10711 feature MANSC domains. Structural alignment of pKP177L with either protein revealed that the region of homology was located in the extracellular domain of pKP177L and the MANSC domain of each protein. The hit with the highest TM-score and the lowest RMSD value for the Georgia strain protein was MANSC Domain-Containing Protein 4 (AF-D4ADQ5-F1-model_v4), a membrane protein from *Rattus norvegicus*. Once again, the region of homology corresponded to the MANSC domain. Sequence alignment using Clustal Omega MSA indicated a sequence identity of 27.42% in this region, although pKP177L contains only three of the seven conserved cysteine residues typically found in the MANSC domain. Indicating pKP177L contains an extracellular domain that is structurally similar to the MANCS domain.

#### *EP153R* (pEP153R)

A Foldseek search using the Ba71v and Georgia strain models of pEP153R identified numerous hits, with e-values ranging from the cut-off threshold of 10⁻³ to 7 × 10⁻⁷. The majority of these hits corresponded to proteins containing a C-type lectin domain (CTLD), with many of the top-ranked hits featuring the NK-type fold of the CTLD.

To explore this further, the amino acid sequence of pEP153R was aligned with that of CD69, a protein identified in Foldseek searches for both Ba71v and Georgia pEP153R and known to contain the typical NK-type CTLD. The alignment demonstrated that pEP153R retains one of the two conserved cysteine residues commonly found in CTLDs, as well as the cysteine residue characteristic of the NK-type CTLD fold (Figure 1). Moreover, five of the seven residues implicated in dimerisation were also conserved.

**Figure 1:**
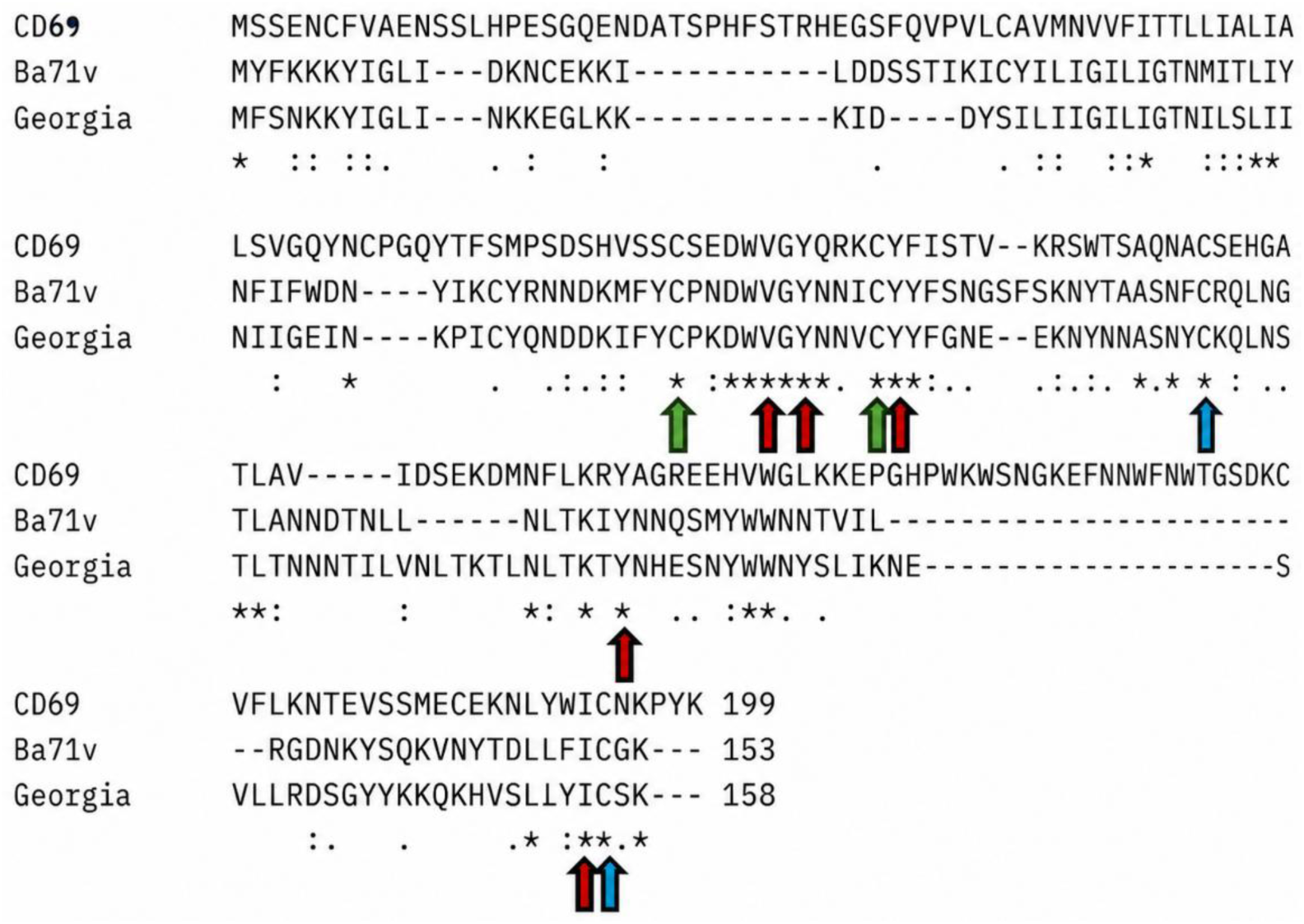
EP153R Sequence alignment. Sequence alignment of CD96 with EP153R Georgia and Ba71v strain sequences. Arrows: Green – NK conserved cysteine residue, blue – CTLD conserved cysteine, red – conserved dimerisation residues. Generated using Clustal Omega web server (https://www.ebi.ac.uk/jdispatcher/msa/clustalo).

To identify the closest homologues from the Foldseek search, a phylogenetic tree was constructed using 16 sequences of pEP153R, including both genotype I and II, alongside the top five Foldseek hits ranked by e-value (Figure 2). The tree revealed that the pEP153R sequences form a distinct cluster, whilst the Foldseek hits form another. The closest related protein was Lectin-like transcript 1 (LLT1), being the only protein to share the most recent common ancestor. Notably, LLT1 was the top hit in both Foldseek searches, producing the highest TM-score, the lowest RMSD, and the most significant e-value.

**Figure 2:**
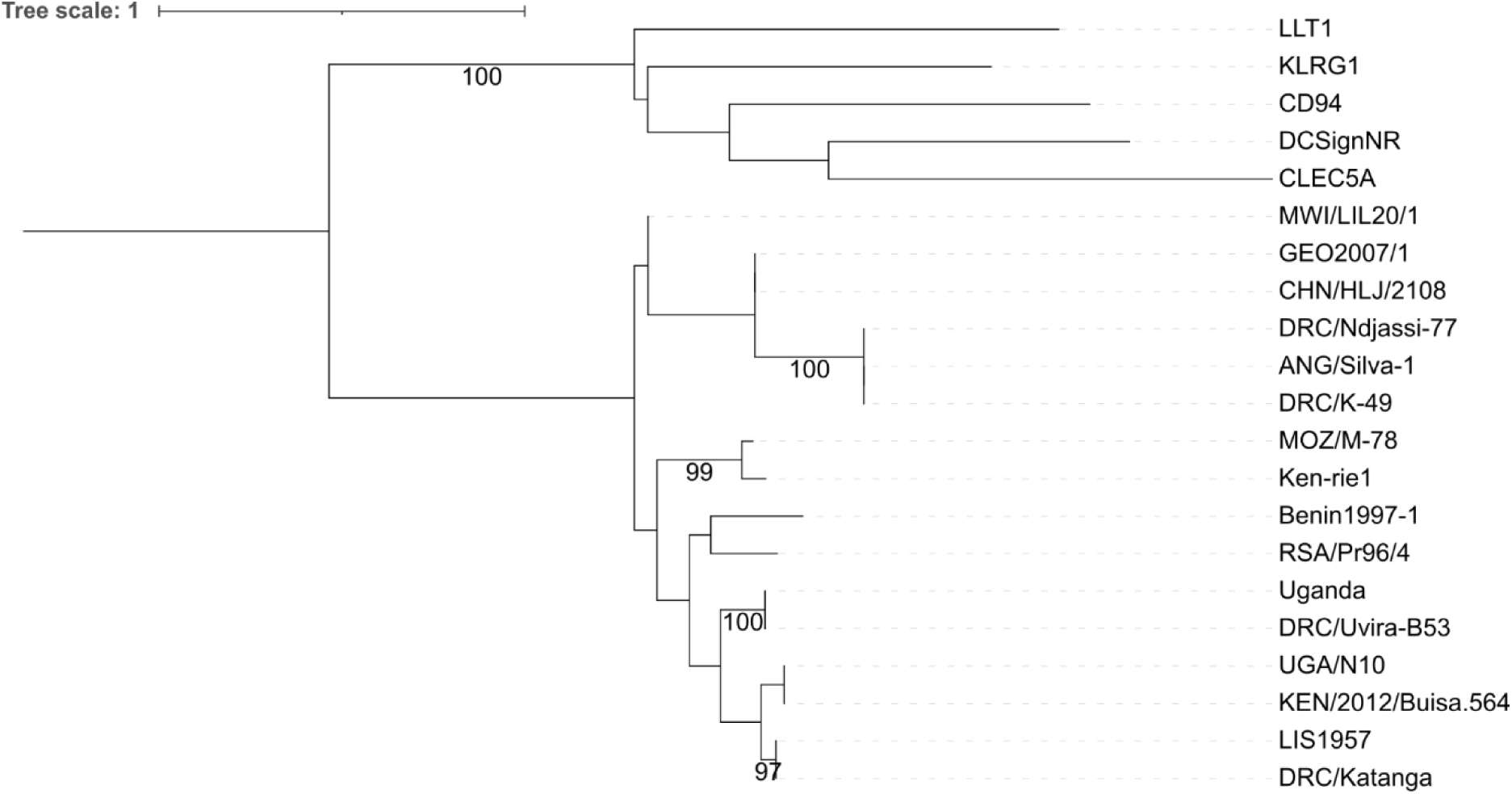
Genetic relatedness of African swine fever virus pEP153R and cellular protein sequences inferred using the maximum likelihood method. Phylogenetic trees were generated using IQ-TREE 3. The percentage of replicate trees in which the associated taxa clustered together in the ultrafast bootstrap test (10,000 replicates) in at least 95% of the trees are shown next to the branches. The VT+G4 model was selected using ModelFinder and the tree was rooted at the mid-point.

Given LLT1 was the closest structural hit to pEP153R, it’s potential to bind LLT1s binding partner, natural killer cell receptor protein 1 (NKR-P1), was assessed by modelling the potential protein-protein interactions in AlphaFold V3. The resulting model presented a similar binding site location and architecture to the canonical LLT1-NKR-P1 interaction when compared to the crystal structure obtained via X-ray diffraction (Blaha *et al*., 2022)(Figure 3)

**Figure 3:**
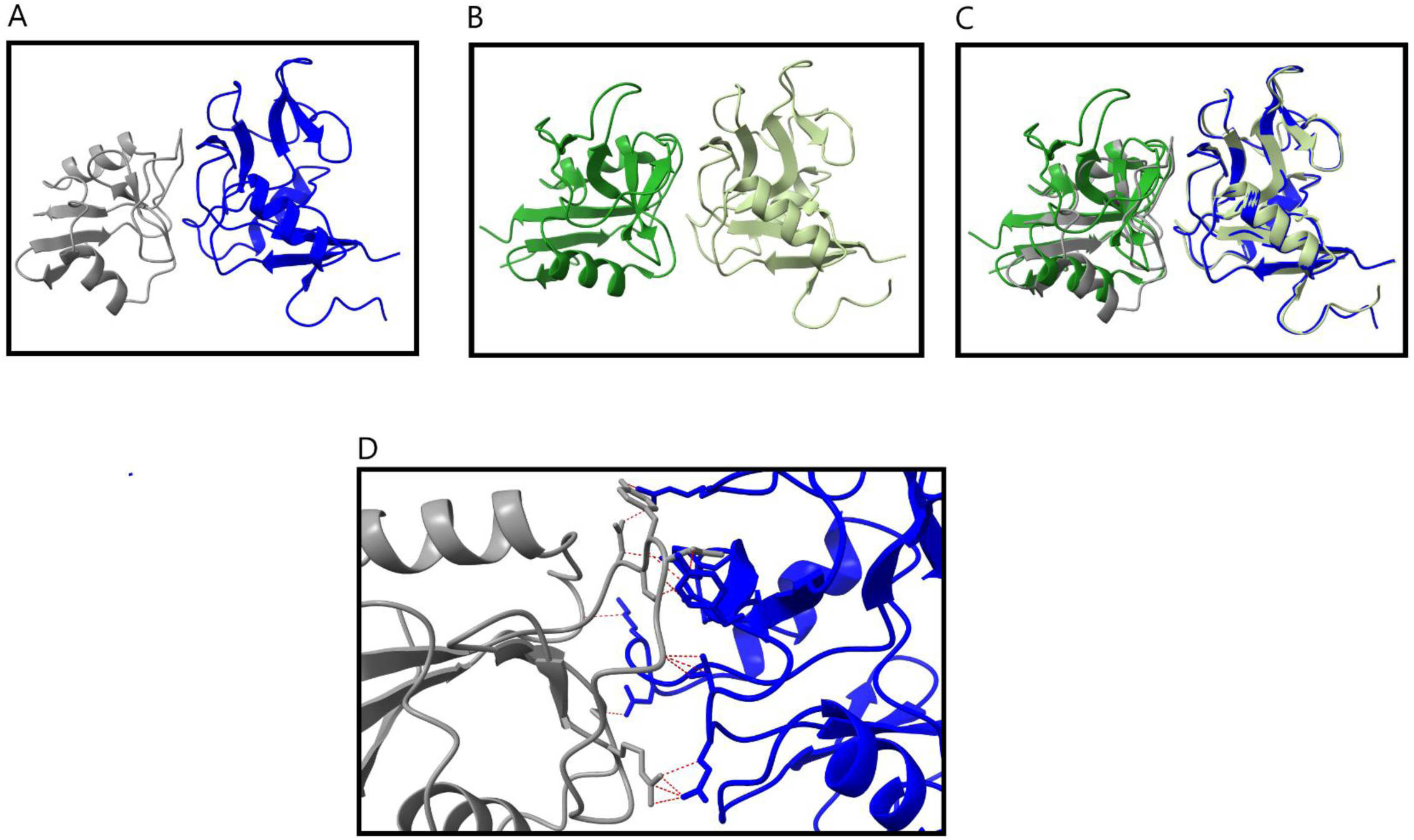
Comparison of pEP153R:LLT1 interaction with NKR-P1:LLT1 interaction. **A.** AlphaFold model prediction of human NKR-P1 (Left) interacting with pEP153R Ba71v (Right). **B.** Crystal structure of human NKR-P1 (Left): LLT1 (Right) receptor: ligand complex. **C.** Overlayed image of A and B that were structurally aligned in ChimeraX v1.6.1. **D.** Zoomed in image showing binding interface from A with contact residues of up to 3Å shown by red dotted lines.

Further analysis in ChimeraX revealed multiple contact residues within a distance of 3Å distributed throughout the binding interface (Figure 3).

### Grouping of candidate proteins for PV generation

Based on the combined data from the *in silico* analysis, proteins were grouped into sets for PV generation, such as their expected presence on the outer membrane or their function, as shown in Table 1 The groupings were designed to encompass all the candidate proteins at least once, while also grouping together the most likely proteins to function in combination.

**Table 1:**
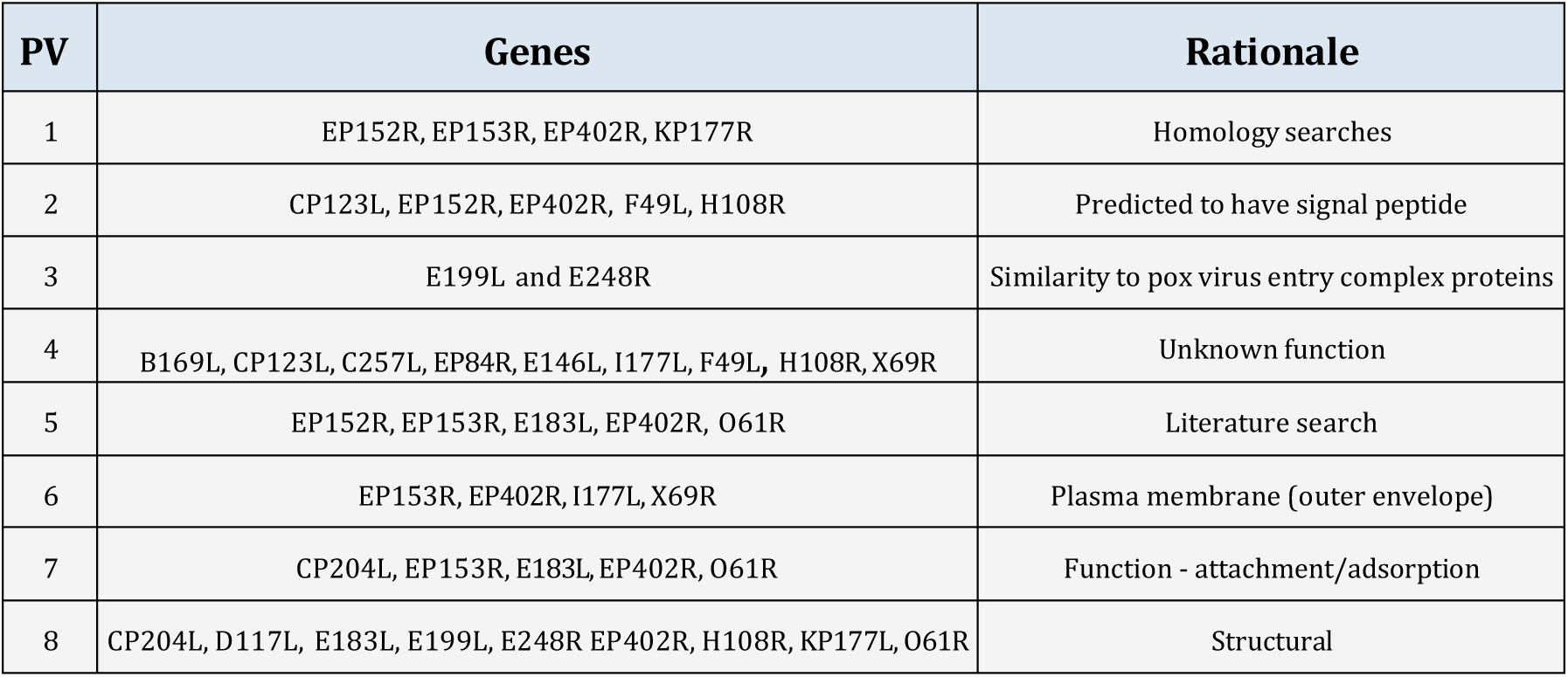
Gene groupings for PV generation. From left to right: PV gene grouping number, the genes within that construct, a short description of the rational for the grouping.

### Generation of rVSV-based ASF-PV and cell line selection

Based on these groupings we next generated PV for each group of proteins. As ASFV proteins in group 7 had already been reported to be involved in attachment or entry, PV7 was used as an initial rVSV-PV to test in various cell lines. rVSV-PV were generated as described in the methods and titrated on four commonly used cell lines (Huh7, Vero, HEK and BHK). Titres ranged from 1.9 × 10^6^ RLU/mL (BHK) to 7.8 × 10^6^ RLU/mL (Huh7) (Figure 4A) with the titres achieved using Huh7 cells being significantly higher than titres achieved from BHK cell (p<0.05).

**Figure 4:**
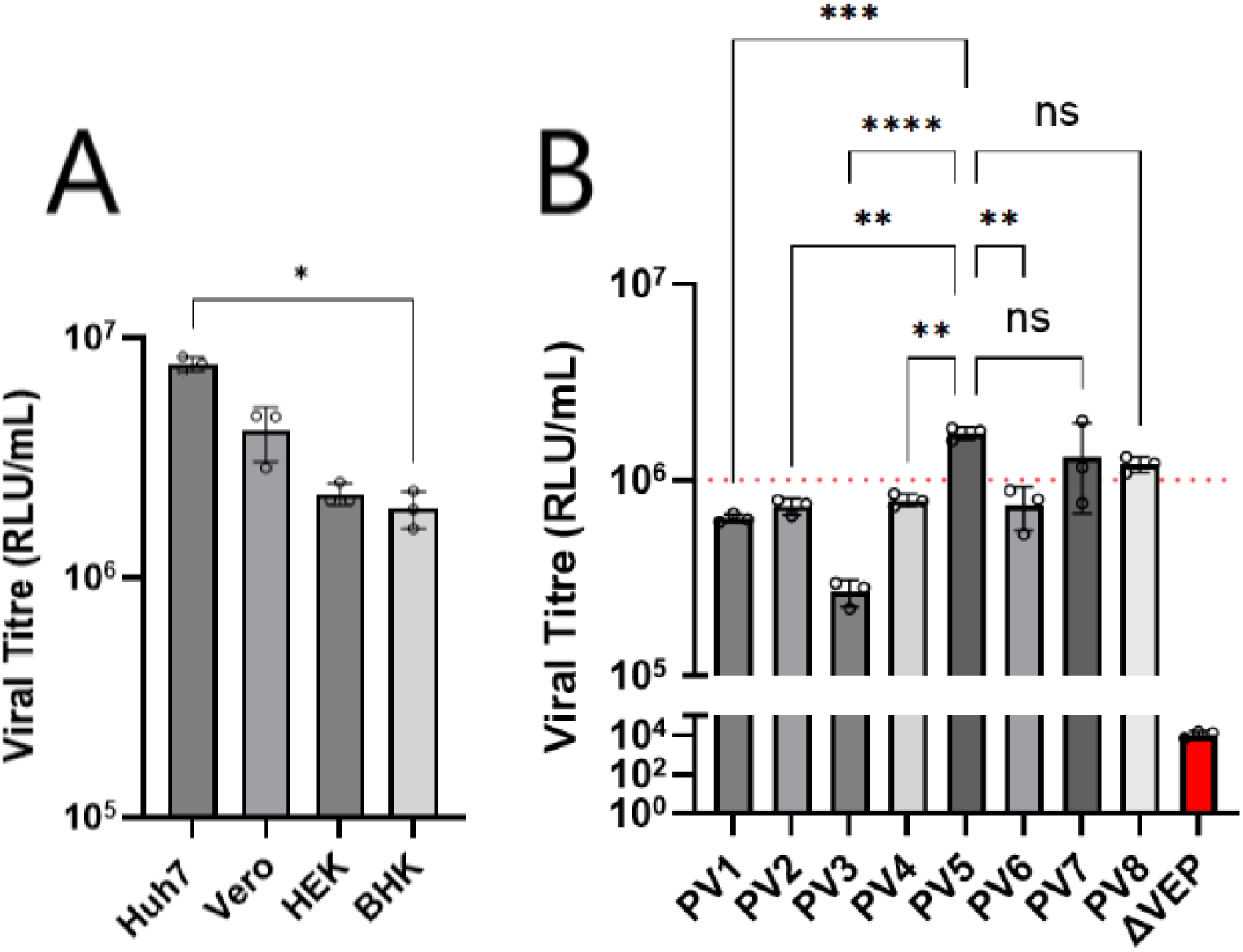
rVSV-PV7 titrated on various cell types and rVSV-PV-1 to 8. **A.** PV titres (RLU/mL) of rVSV-PV7 (10 cm dish) on Huh7, Vero, HEK293T/17 and BHK cells. **B.** PV titres of rVSV PV1-8 (6-well plate) on Huh7 cells. Red dotted line indicates 1x10^6^ cut off for functional titre. Error bars display standard deviation. N=3 biological repeats with each run as 2 technicals. Normality was established via a Shapiro-Wilk test and statistical differences between titres were established via a Brown-Forsythe test and one-way ANOVA with Tukey’s multiple comparison. * = p < 0.05; ** = p < 0.01; *** = p < 0.001; **** = p < 0.0001; NS = p ≥ 0.05.

Next, all eight of the PV constructs were generated for the Ba71v strain. These rVSV-ASF-PV (Ba71v) were then titrated on Huh7 cells. Titres ranged from 2.5 × 10^5^ to 1.7 × 10^6^ RLU/mL. PV1, 2, 3, 4, and 6 generated titres under the 10^6^ threshold for functional PV with titres of 6.5, 7.6, 2.5, 7.9 and 7 × 10^5^ RLU/mL respectively. Whilst PV5, 7 and 8 generated titres above this threshold, with titres of 1.7, 1.3 and 1 × 10^6^ respectively (Figure 4B). PV5 therefore generated the highest titre which was significantly higher than the titres achieved by PV1, 2, 3 or 4 (p values: PV1 = 0.0009, PV2 = 0.0024, PV3 = <0.0001, PV4 = 0.004). However, there was no statistically significant difference when compared to PV6, 7 or 8 (p-values: >0.05).

In examining the three PV constructs that generated titres above the 10⁶ RLU/mL threshold (PV5, PV7 and PV8), it became apparent that all three groups shared a common subset of three ASFV candidate proteins (pO61R, pE183L, pEP402R). These shared proteins were therefore selected for further comparative analysis.

The three proteins were assessed in pairwise combinations and compared to PV5, to ascertain which proteins were required to produce the highest titres. The combination of these three candidate proteins (PV9) produced a titre of 1.5 × 10^7^ RLU/mL, an 8.8 fold increase, that was significantly higher (p value= 0.0001) than the 1.7 × 10^6^ RLU/mL produced by PV5. All three of the PV displaying pairs of the three proteins (EP402R+O61R, EP402+E183L and O61R+E183L) generate higher titres than PV5; 7.6, 7.5 and 4.7 × 10^6^ RLU/mL respectively. Moreover, PV9 titres were at least a 2-fold increase from any paired combination PV, and significantly higher with p values of <0.0001 for EP402+O61R, EP402+E183L and O61R+E183L (Figure 5).

**Figure 5:**
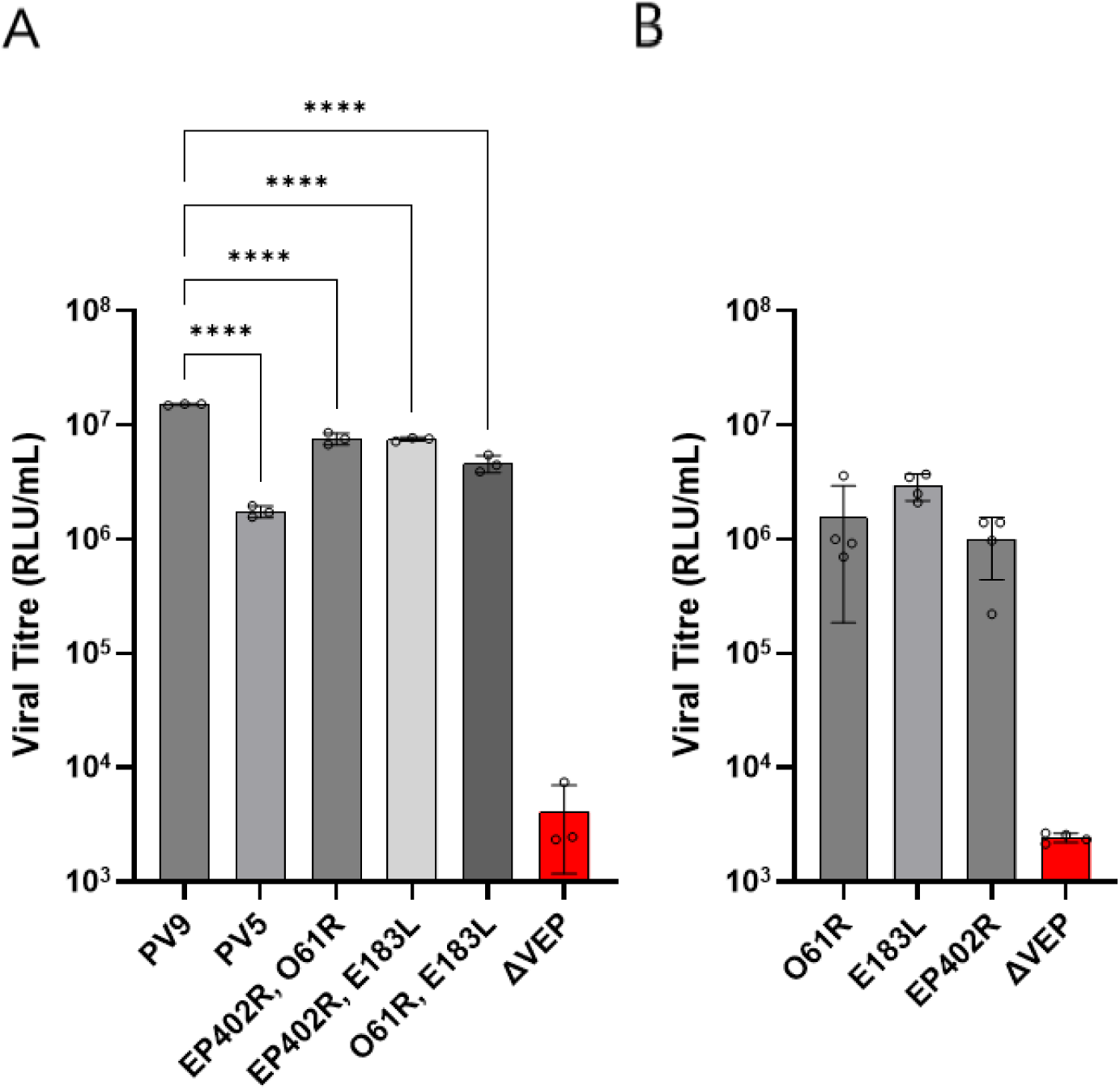
Titres for rVSV-PV9 and its combinations. **A.** rVSV-PV titres (RLU/mL) of PV9 (pEP402R, pE183L and pO61R), PV5 (pEP402R, pE183L, pEP152R, pEP153R and pO61R), paired combinations of PV9 proteins and ΔVEP control. Normality was established via a Shapiro-Wilk test and statistical differences between titres were established via a Brown-Forsythe test and one-way ANOVA with Tukey’s multiple comparison. **B.** rVSV-PV titres (RLU/mL) of O61R, E183L, EP402R and ΔVEP control. proteins Normality was established via a Shapiro-Wilk test and statistical differences via Kruskal-Wallis test was used followed by Dunn’s multiple comparisons test. * = p < 0.05; ** = p < 0.01; *** = p < 0.001; **** = p < 0.0001; NS = p ≥ 0.05.

### Investigating rVSV-ASF-PV9 cell tropism

To investigate the cell tropism of the rVSV-ASF-PV9, both the Ba71v and Georgia PV were titrated onto various cell lines known to be permissive to authentic ASFV. This included Vero cells, the cell line Ba71v is adapted to replicate in, PLTA58s, an immortalised macrophages-like cell line that express pig macrophage markers, and porcine alveolar macrophages (PAMs), a primary cell culture. Huh7 cells, derived from a human hepatocellular carcinoma, were included as they were the most permissive for the PV.

Ba71v and Georgia PV produced titres of 4.27 × 10^7^ and 2.39 × 10^7^ RLU/mL respectively, and there was no statistical significance between the PV titres from Huh7 cells. Titrations using Vero cells produced titres of 3.14 × 10^6^ for Ba71v and 1.24 × 10^6^ RLU/mL for the Georgia PV, Georgia PV titres where significantly lower than Ba71v (p = 0.0283). Titrations using PLTA58 cells produced titres of 1.83 × 10^6^ and 1.55 × 10^6^ RLU/mL with no statistical significance between titres. Unfortunately, the titrations using PAMs did not generate raw RLU values higher than the cell only background and thus, did not generate a titre, nor was there a rise in ΔVEP control which was predicted to be higher due to constitutive activation of macropinocytosis in macrophages (Figure 6).

**Figure 6:**
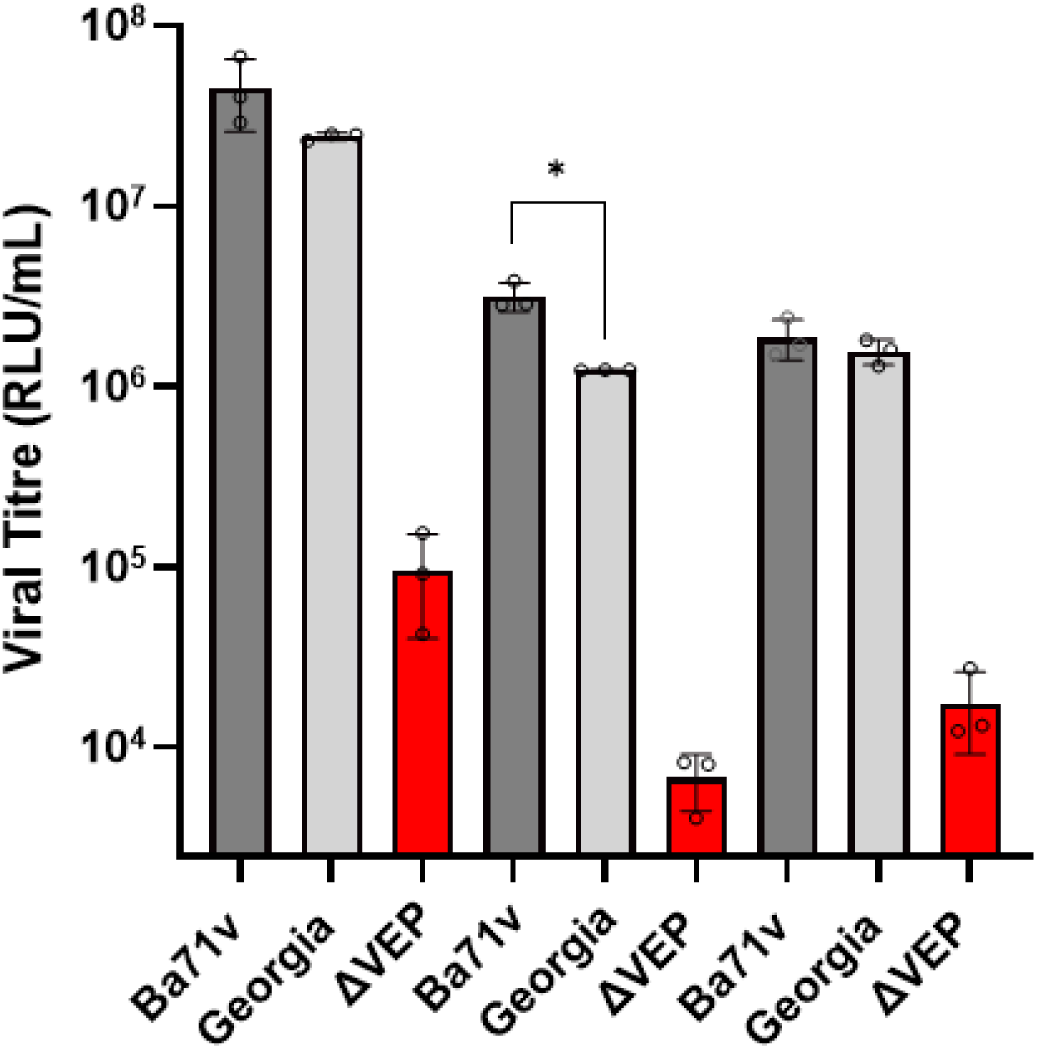
rVSV-ASFV-PV9 (Ba71v and Georgia) titrated on multiple cell types. rVSV-ASFV-PV9 (Ba71v and Georgia) titres (RLU/mL) in multiple cell types. Dark grey bars show Ba71v, light grey Georgia strains and red the negative control (ΔVEP for respective cell lines). Normality was established via a Shapiro-Wilk test and statistical differences between titres were established via a Brown-Forsythe test and one-way ANOVA with Tukey’s multiple comparison. Error bars display stand deviation. N=3 biological repeats with each run as 2 technicals. * = p < 0.05; ** = p < 0.01; *** = p < 0.001; **** = p < 0.0001; NS = p ≥ 0.05.

### Investigating pEP153R as a cell adhesion protein

The results of the bioinformatic analysis of pEP153R, reported earlier, indicated that pEP153R might act as a cell adhesion protein and therefore aid viral cell entry. To test this theory rVSV-ASF-PV9 was generated to also include pEP153R for both the Ba71v and Georgia strains. rVSV-ASF Ba71v and Georgia both with and without pEP153R were then titrated onto Huh7, Vero and PLTA58 cell lines. The only significant difference (p Value >0.05) between PV constructs with and without pEP153R was for the Georgia strain PV titrated on Huh7 cells (Figure 7).

**Figure 7:**
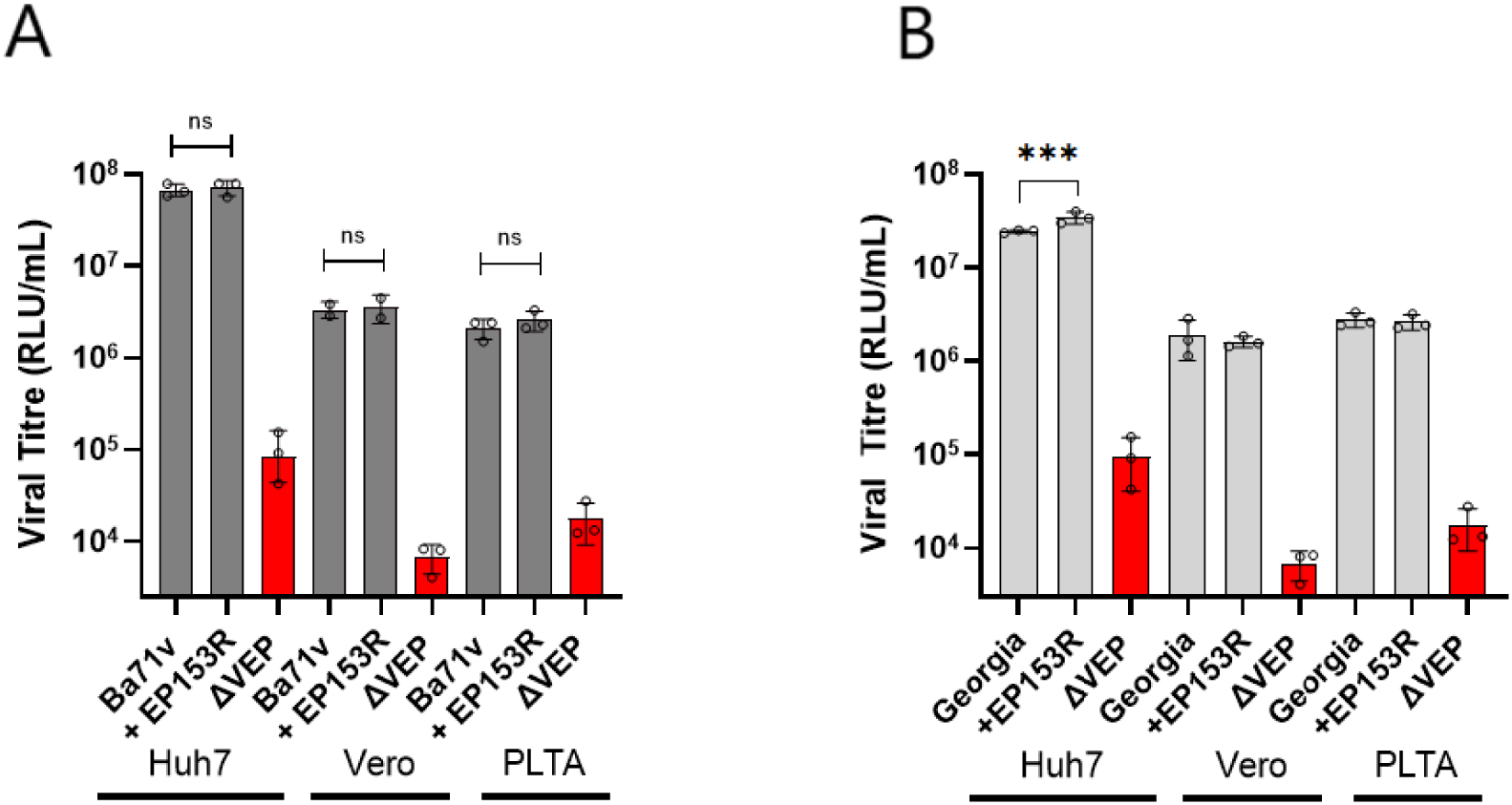
rVSV-ASF-PV9 Ba71v and Georgia, with and without pEP153R, titrated on multiple cell lines. **A.** rVSV-ASF-PV9 (Ba71v) titres (RLU/mL) with and without pEP153R. **B.** rVSV ASF PV9 (Georgia) titres (RLU/mL) with and without pEP153R. Dark grey bars show Ba71v and grey Georgia strains. Red columns show the respective ΔVEP control. Normality was established via a Shapiro-Wilk test and statistical differences between titres were established via a Brown-Forsythe test and one-way ANOVA with Tukey’s multiple comparison. Error bars display stand deviation. N=3 biological repeats with each run as 2 technicals. * = p < 0.05; ** = p < 0.01; *** = p < 0.001; **** = p < 0.0001; NS = p ≥ 0.05.

## Discussion

This study aimed to utilise viral pseudotype technology to identify key ASFV proteins involved in cell entry. To our knowledge, this approach has not previously been applied to identify VEP. As higher PV titres correspond to increased levels of PV transduction, we sought to determine whether a systematic and rational approach could be applied to identify the glycoproteins required for ASFV viral entry. Ultimately, this work aimed to establish an ASF-PV system capable of mimicking viral entry under reduced containment conditions.

A panel of 19 candidate ASFV proteins was selected based on published literature, bioinformatic analyses and existing knowledge of viral envelope proteins. Due to the large number of candidates identified, these proteins were organised into eight groups according to predicted or reported involvement in viral entry, structural similarity to envelope proteins from other viruses, or other functional characteristics. This approach enabled multiple hypotheses regarding ASFV entry mechanisms to be investigated in parallel whilst ensuring that all candidate proteins were assessed at least once.

Application of the VSV pseudotype system incorporating ASFV candidate proteins generated multiple functional PV titres. Comparative analysis demonstrated that the three highest-titre constructs shared a common subset of ASFV proteins: O61R, E183L and EP402R. Subsequent deconstruction experiments confirmed that the combination of these three proteins alone (PV9) produced titres an order of magnitude higher than any other construct tested. Importantly, PV9 titres were significantly greater than those produced by any pairwise combination, indicating that all three proteins are required for optimal entry. Moreover, PV9 (Ba71v) titres were only significantly different to PV9 (Georgia) when titrated on Vero cells, which Ba71v is adapted to, showing this ASF PV system shows strain specific tropism consistent with authentic virus.

The requirement for multiple ASFV proteins to achieve efficient PV entry is consistent with the complex entry mechanisms reported for other large, enveloped DNA viruses. ASFV has previously been proposed to utilise an entry fusion complex (Urquiza *et al*., 2024, Matamoros *et al*., 2020), although direct evidence remains limited. Several ASFV proteins exhibit structural similarity to components of the poxvirus fusion entry complex. pE199L shares similarities with three of the eleven proteins comprising the poxvirus entry fusion complex, namely A16, G9 and J5 (Iyer *et al*., 2001), whilst pE248R shares amino acid sequence similarity with the vaccinia virus L1 protein (Hernaez *et al*., 2016). Our findings may therefore support the hypothesis that ASFV entry involves coordinated multi-protein interactions. However, the proteins identified within PV9 do not display structural similarity to known components of the poxvirus entry fusion complex. Although the precise mechanistic roles of pO61R, pE183L and pEP402R during ASFV entry remain unclear, this study provides novel functional evidence supporting their combined involvement in the entry process and establishes a foundation for future work investigating ASFV entry mechanisms using PV systems.

The rVSV-ASF-PV9 pseudotype demonstrated tropism consistent with authentic ASFV and produced functional titres in both Vero and PLTA58 cells, which are known to support ASFV replication. However, no detectable signal was observed in primary PAMs. The inability to generate measurable titres in PAMs may reflect limitations of the assay system rather than a true restriction in viral entry, particularly given the high background activity associated with constitutive macropinocytosis in macrophages. As the assay depends upon delivery of the viral core into the cell followed by luciferase gene expression, it is possible that pseudotyped particles are internalised by macrophages but subsequently fail to escape the endosomal compartment prior to lysosomal fusion, resulting in degradation of the PV before reporter expression can occur. Consequently, no luciferase signal would be detected. This highlights a potential limitation of PV systems as surrogates for viruses that preferentially target highly phagocytic cell types and represents an area requiring further investigation. Alternative culture systems, including blood monocyte-derived and bone marrow-derived macrophages, are known to support replication of ASFV field isolates and may therefore provide more physiologically relevant models for future PV studies (Goatley *et al*., 2022, Rathakrishnan *et al*., 2022). Moreover, other macrophage-like lines have also been shown to support ASFV replication and could also be explored as alternatives (Masujin *et al*., 2021, Portugal *et al*., 2020).

As bioinformatic analysis suggested pEP153R has structural similarity to LLT1, which can bind glycosaminoglycans via its lectin domain (Gange et al., 2004). This raised the possibility that pEP153R may similarly interact with glycosaminoglycans on host cells and thereby contribute to viral attachment or adhesion. To investigate whether pEP153R plays a role in cell adhesion, rVSV-ASF-PV9 were generated both with and without pEP153R on their surface, and their capacity to transduce a range of cell types was subsequently compared. Inclusion of pEP153R did not significantly alter PV transduction across multiple cell types, suggesting either that pEP153R does not bind glycosaminoglycans or that such binding does not substantially influence cell adhesion within this system. However, pEP153R has previously been implicated in immune evasion, with studies demonstrating that it reduces surface expression of MHC-I by inhibiting the exocytosis of newly synthesised molecules to the plasma membrane (Hurtado *et al*., 2011). This activity would limit antigen presentation and thereby reduce recognition by cytotoxic T cells, supporting viral persistence during infection. In addition to this established mechanism, the presence of a C-type lectin-like domain within pEP153R and its structural similarity to LLT1 raises the possibility that it may also function through receptor mimicry. LLT1 and its receptor, natural killer cell receptor protein 1 (NKR-P1), form clusters proposed to facilitate inhibitory immune synapse formation (Blaha *et al*., 2022). KLRG1, the next closest Foldseek hit Plays an inhibitory role of natural killer and T-cell functions after binding their non-MHC ligands (Li *et al*., 2009). Moreover, CD94, the next closest hit has also been shown to play similar inhibitory roles for natural killer and T-cell functions (Le Drean *et al*., 1998). Thus, pEP153R could mimic these interactions to evade early antiviral responses. Whilst direct evidence for such interactions involving pEP153R is currently lacking, this represents an area that warrants further investigation in future studies but was beyond the scope of this project.

Several limitations of this study should also be acknowledged. The PV system models viral entry in isolation and therefore may not mimic the entire process of authentic ASFV entry. Furthermore, the density, conformation and copies of ASFV proteins incorporated into VSV particles may differ from those present on native virions. Consequently, additional ASFV proteins or host factors not identified in this study may also contribute to authentic ASFV entry *in vivo*.

In summary, this study establishes a novel ASFV pseudotype system, as a low-containment authentic virus surrogate and identifies pO61R, pE183L and pEP402R as key contributors to efficient viral entry. The requirement for multiple ASFV proteins supports the concept that ASFV entry is mediated through coordinated multi-protein interactions rather than a single viral attachment factor. Beyond improving understanding of ASFV biology, the development of a functional pseudotype system may provide a valuable tool for future studies investigating viral tropism, host–virus interactions, and entry-targeted antiviral or vaccine strategies under reduced biocontainment conditions.

## Methods and materials

### Bioinformatics

Gene sequences were obtained from the GenBank database for both Ba71v and Georgia 2007 strains. Sequences were scanned against the InterPro protein signature databases using the InterProScan tool (Jones *et al*., 2014). Signal peptide predictions were carried out by SignalIP v6.0 (Teufel *et al*., 2022) using the following settings: organism: other, output: long output, model: slow.

3D structural predictions were performed by uploading amino acid sequences to ColabFold AlphaFold V3 server (https://search.foldseek.com/search) (Abramson *et al*., 2024) with default parameters. PDB files obtained were uploaded to the Foldseek server (https://search.foldseek.com/search, version 10-941cd33) and a standard search run to search for structural homology. The PDB files were processed using icn3D (Wang *et al*., 2020) for 2D images and ChimeraX v1.6.1 (Meng *et al*., 2023) for structural analysis.

Sequences (Supplementary Table 2) were aligned using clustal omega (v1.2.4) and phylogeny performed using iq-Tree (v3.0.1) (Minh *et al*., 2020) using Modelfinder (Kalyaanamoorthy *et al*., 2017) and ultrafast bootstrapping (Hoang *et al*., 2018). Trees were displayed using the interactive tree of life (Letunic and Bork, 2024).

### Mammalian cell lines

Cell lines (BHK-21 (CCL-10), Huh7 (JCRB0403), HEK 293T/17 (ACS-4500), VeroE6 (CRL-1586)) were maintained in a humidified atmosphere containing 5% CO2, at 37 °C, in Dulbecco’s Modified Eagle’s Medium-High Glucose (DMEM; Sigma, UK), containing either 5 (BHK-21 only) or 10% foetal bovine serum (FBS; Sigma, UK). Cells were cultured in either 10cm TPP tissue culture petri dishes or 10cm Nunc cell culture petri dishes (HEK293T only). All cell lines were passaged three times per week. The PLTA58 macrophage-like cell line and primary porcine macrophages derived from bronchoalveolar lung wash (Goatley *et al*., 2022) were provided by the African swine fever vaccinology group, The Pirbright Institute.

### ASFV gene construct design and synthesis

The DNA sequence for each ASFV gene was codon optimised by GeneArt (Thermofisher). The sequences each comprised the following, from 5’ to 3’: 6bp random bases (GATCTA) to enable restriction enzyme binding, *Nhe*I restriction enzyme site (GCTAGC), a Kozak sequence (GCCACC) to improve gene transcription, ASFV human codon optimised gene sequence, haemagglutinin tag (TACCCCTACGACGTGCCCGACTATGCC), stop codon (TGA), *Apa* I restriction enzyme site (GGGCCC), 6bp random sequence to enable efficient RE activity during cloning (GATCTA). Sequences were uploaded to and ordered from Eurofins genomics. All genes were cloned into pCDNA3.1 expression vectors and sequences confirmed by sanger sequencing (Eurofins, LightRun).

### Production of rVSV based pseudotyped viruses

First, 4 × 10^5^ HEK293T/17 cells were seeded per well of 6-well tissue culture plate or 5 × 10^6^ for a 10 cm tissue culture dish and incubated overnight. Then, the media was replaced with fresh DMEM + 10% FBS (Sigma, UK), and the cells transfected with various quantities of each VEP plasmid DNA using PEI (Sigma, UK). The following day the media was replaced, and cells were infected with stocks of amplified rVSV-Luciferase at an MOI of 0.5. These were then incubated at 37°C, 5% CO_2_ for one hour before the media was removed and the cells washed with PBS (Sigma, UK). Fresh media was added to each well and the cells incubated at 37 °C, 5% CO_2_. The supernatant was harvested, and syringe filtered through a 0.45 μM filter (Sigma, UK) at 48 hours post-infection.

### Titration of rVSV-ASF pseudotyped viruses

Pseudotyped virus was serially diluted 1:2 for at least 3 dilutions, in duplicate, across a Nunc^TM^ F96 LumiPlate. Then, 2 × 10^4^ cells were added to each well, including a set of cell-only control wells. The plate was then incubated at 37 °C, in 5% CO_2_ for 48 hours. To prepare the plate to be read by a Glo-max^®^ Discover microplate reader (Promega, UK), the media was removed and 15 μL of Bright-Glo^®^ reagent and 15 μL of PBS (Sigma, UK), added to each well. The plate was placed on a plate shaker for 2 minutes at 1,000 rpm and the luminescence read 5 minutes. Titres are expressed as average RLU/mL of replicates.

### Statistics

Normality was assessed using the Shapiro–Wilk test. Equality of variances was evaluated using the Brown–Forsythe test. Where data were normally distributed and variances were equal, statistical differences between titres were analysed using one-way ANOVA with Tukey’s multiple comparisons test. Where variances were unequal, Welch’s ANOVA was performed followed by Dunnett’s T3 multiple comparisons test. Where normality was not established a Kruskal-Wallis test was used followed by Dunn’s multiple comparisons test.

When comparing two titres only, normality was assessed using the Shapiro–Wilk test. Equality of variances was evaluated using an F-test. For normally distributed data with equal variances, differences between titres were analysed using an unpaired Student’s t-test. Where variances were unequal, Welch’s t-test was applied.

## CRediT author contributions

GLH – Conceptualisation, data curation, formal analysis, investigation, writing – original draft.

CLN – Conceptualisation, funding acquisition, investigation, resources, supervision, writing – reviewing and editing

EW – Conceptualisation, funding acquisition, investigation, supervision, writing – reviewing and editing

## Acknowledgements

We are grateful to Tom Foster for his early insights and discussions that helped shape this study.

## Funding

GLH was supported by a studentship awarded by the Pathogens and Host Defences doctoral training programme (Sussex/Pirbright/Surrey/DSTL). CLN was supported by UKRI grants [BBS/E/I/00007034](https://www.sciencedirect.com/unsupported_browser#gs2), [BBS/E/I/00007037](https://www.sciencedirect.com/unsupported_browser#gs2), [BBS/E/PI/0000230002A](https://www.sciencedirect.com/unsupported_browser#gs2) and [BBS/E/PI/000023NB0004](https://www.sciencedirect.com/unsupported_browser#gs2).

## Notes

### Competing Interest Statement

The authors have declared no competing interest.

